# The interplay between temperature and growth phase shapes the transcriptional landscape of *Pseudomonas aeruginosa*

**DOI:** 10.1101/2025.08.28.672984

**Authors:** Rachel E. Robinson, Michael J. Gebhardt, Joanna B. Goldberg

## Abstract

*Pseudomonas aeruginosa* is a highly versatile bacterium capable of surviving and often thriving in stressful environmental conditions. Here we report the effect of two environmental conditions, temperature and growth phase, on the *P. aeruginosa* PAO1 transcriptome. As *P. aeruginosa* is well-known for its growth phase dependent phenotypes and gene regulation, our goal was to determine how temperature altered global gene expression at exponential versus stationary phase and to characterize how growth phase affects thermoregulation. To do this, we grew PAO1 in parallel at 25°C and 37°C and sampled the same populations first at exponential phase and then again at stationary phase and assessed gene expression by RNA-sequencing. We found that temperature regulated hundreds of genes at, and unique to, exponential and stationary phase. We also grew PAO1 and an isogenic Δ*lasR* mutant at 25°C and 37°C and sampled populations at stationary phase to define LasR-regulated genes at each temperature by RNA-sequencing. LasR regulated most of its target genes similarly at 25°C and 37°C, although we identified a subset of genes whose regulation by LasR was affected by temperature. This work provides a comprehensive thermoregulon for PAO1 at two distinct growth phases, as well as growth phase transcriptomics at two temperatures, and expands our understanding of quorum sensing regulation under different environmental conditions that *P. aeruginosa* encounters.

**IMPORTANCE:** *Pseudomonas aeruginosa* is a highly adaptable opportunistic pathogen with a repertoire of mechanisms for surviving in diverse and often challenging environments yet is most often studied at 37°C as the optimum temperature for growth. To better understand how this bacterium survives in the environment versus the human body, we performed transcriptomics on *P. aeruginosa* grown at 25°C and 37°C. At each temperature, we examined both exponential and stationary phases. We also determined the LasRI quorum sensing regulon at 37°C compared to 25°C using a Δ*lasR* mutant, which uncovered a suite of previously unrecognized LasR-regulated genes. Our work provides a comprehensive transcriptomic resource for thermoregulation of *P. aeruginosa* at two growth phases, as well as growth phase and LasR regulation at two temperatures.

## INTRODUCTION

*Pseudomonas aeruginosa* is a versatile opportunistic bacterial pathogen that causes burn, wound, corneal, and respiratory infections in immunocompromised patients (1). It is a particular health burden for people with the genetic disorder cystic fibrosis (CF), in whom *P. aeruginosa* can cause chronic and often life-long infections that result in significant morbidity and mortality (2). Antibiotic resistance further underscores *P. aeruginosa* as a major public health concern, with the emergence of untreatable strains resistant to last line antibiotics (3).

Although its optimum temperature for growth is 37°C, *P. aeruginosa* can survive temperatures from 4°C to as high at 42°C – a wide range that distinguishes it from other *Pseudomonads* (4, 5). Accordingly, it can be found in a wide range of environments, although it is more often isolated from anthropogenic locations such as sinks, hospital surfaces, and medical devices like ventilators and catheters (6). The transition from a contaminated surface to the human host inherently involves a change from ambient temperature to human body temperature, 37°C (or higher in cases of fever). However, most laboratory studies of *P. aeruginosa* physiology and pathogenesis have historically been conducted with bacterial cells grown at 37°C; this is despite the intrinsic relevance of temperature changes to nosocomial infections in which *P. aeruginosa* transitions from ambient or room temperature to human body temperature. As many *P. aeruginosa* infections are acquired from healthcare settings, investigating how this bacterium adapts to different temperatures associated with nosocomial infections will provide insights into mechanisms that are important for its pathogenesis.

It is commonly appreciated that many bacterial pathogens sense and respond to human body temperature by regulating the expression of virulence factors (7, 8). More recently, the mechanistic basis for the thermoregulation of a few virulence factors in *P. aeruginosa* specifically has been studied (9–13). Likewise, a few transcriptomic studies have shown how *P. aeruginosa* responds to growing at ambient (22°C or 28°C) versus human body (37°C) temperature, although these studies were limited to examining global gene expression at a single point in the growth curve (i.e., stationary phase) (14, 15). Extensive and complex transcriptional rewiring occurs in *P. aeruginosa* populations at stationary phase (16–19), largely due to the multiple quorum sensing systems active at this growth phase. Quorum sensing is a system for sensing the density of kin cells and subsequently regulating many genes, including virulence factors and secreted products (reviewed in 16, 17). In the *P. aeruginosa* lab strain PAO1, LasRI is the master quorum sensing system. In brief, LasI synthesizes the diffusible homoserine lactone (HSL) autoinducer molecule 3-oxo-C12, which complexes with and activates the transcriptional activator LasR at higher cell densities. Given the importance of temperature to the various stages of *P. aeruginosa* as an opportunistic pathogen, we wished to investigate global transcriptional thermoregulation at both exponential and stationary phases, as well as to compare how growth at different temperatures impacts the effect of growth phase on the global transcriptome.

Here, we used RNA-sequencing to determine how *P. aeruginosa* PAO1 adapts to growth at an ambient temperature of 25°C versus human body temperature of 37°C, at both exponential and stationary phases. This also allowed us to compare how growth phase affects gene expression in cells grown at 25°C versus 37°C. We further examined how temperature affects regulation by the quorum sensing regulator LasR. These experiments reveal that the thermoregulon in *P. aeruginosa* depends highly on growth phase and that growth phase regulates the majority of the transcriptome similarly at both temperatures tested. We also show that while LasR regulates most target genes to the same degree at both 25°C and 37°C, a few genes were regulated by LasR uniquely in response to different temperatures. By growing strains at 25°C, we identified genes that had not been previously recognized as LasR-regulated in studies conducted at 37°C. This work has expanded our knowledge of how *P. aeruginosa* adapts to growing at two common temperatures in unique ways depending on growth phase.

## RESULTS

### Temperature globally regulates the expression of distinct genes at exponential and stationary phase

To identify genes regulated by temperature at exponential versus stationary phase, PAO1 was grown overnight at 37°C and each of three biological replicates was used to inoculate two ‘paired’ cultures, one of which was incubated at 37°C and the other at 25°C. RNA was extracted from an equal number of cells from each of the cultures first at exponential phase (OD_600_ of 0.5) and then again at stationary phase (OD_600_ of 2.0) as depicted in Fig. 1A and followed by RNA-sequencing. We first examined gene expression at 37°C versus 25°C, hereafter called thermoregulation, at each growth phase by differential expression analysis.

**Figure 1.**
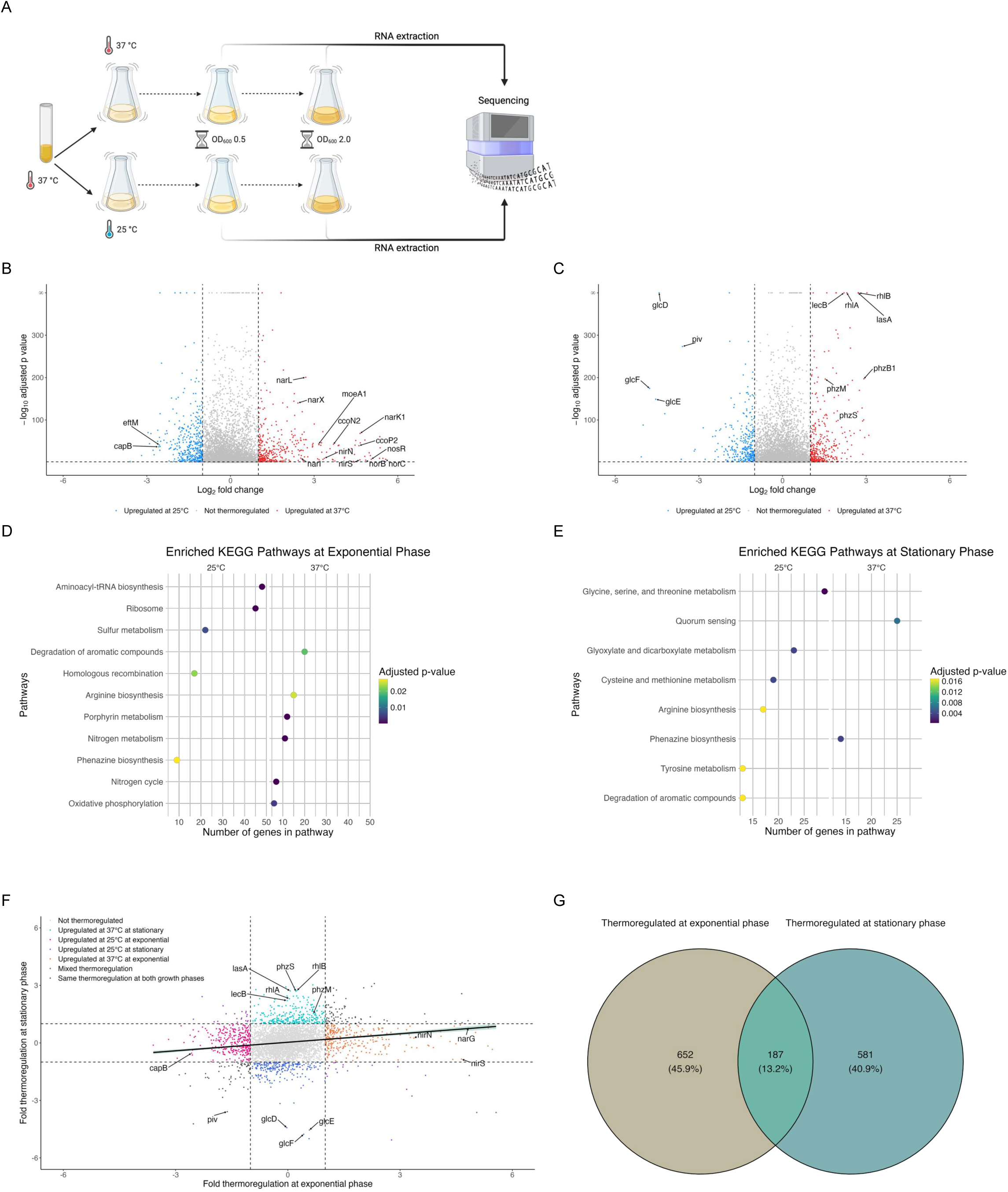
Temperature regulates the expression of hundreds of distinct genes in *P. aeruginosa* at both exponential and stationary phases. A) Diagram of experimental setup for RNA-seq of *P. aeruginosa* PAO1 grown at two temperatures and sampled at two growth phases. PAO1 was grown overnight at 37°C and then subcultured in parallel at 37°C and 25°C. At exponential phase, RNA was extracted from ∼10^9^ cells and the populations allowed to continue growing until stationary phase, when RNA was extracted again from the same number of cells. RNA was then sequenced on an Illumina NextSeq2000. Created with BioRender.com. B, C) Volcano plots showing the thermoregulation of PAO1 transcripts at exponential phase (B) and stationary phase (C). Differential gene expression analysis (DESeq2) was used to compare gene expression at 37°C to 25°C. Transcripts with an absolute value of fold change greater than 2 (vertical dashed lines) and an adjusted p-value < 0.05 (horizontal dashed line) were considered thermoregulated, with transcripts upregulated at 37°C represented by red points and transcripts upregulated at 25°C represented by blue points. Transcripts that did not change by an absolute value of fold change greater than 2 or were not statistically significant (adjusted p-value > 0.05) are depicted in gray. Genes of interest are annotated. D, E) Metabolic pathways significantly enriched (adjusted p-value < 0.05) at exponential phase (D) and stationary phase (E) at 25°C (left panel for each D and E) and 37°C (right panel for each D and E). Enrichment was determined by gene set enrichment analysis using KEGG metabolic pathways for *P. aeruginosa*. F) The fold thermoregulation (expression at 37°C/25°C) of statistically significant transcripts was plotted at exponential phase (x-axis) versus stationary phase (y-axis). Transcripts are colored according to how growth phase affected thermoregulation, with genes of interest annotated. Linear regression with 95% confidence intervals is shown. G) Venn diagram showing at which growth phase (exponential, stationary, or both) statistically significant transcripts were thermoregulated.

### Thermoregulation at exponential phase

At exponential phase, temperature affected the expression of 791 genes by 2-fold or more (adjusted p value < 0.05), indicating that 13.84% of the annotated genome was regulated by temperature (Fig. 1B, with select genes of interest labelled, and Data Set S1). Of these differentially regulated genes, 386 were upregulated at 37°C compared to 25°C and 405 were upregulated at 25°C compared to 37°C. Perhaps unsurprisingly, many thermoregulated genes are involved in metabolism and/or respiration. Many of the biological pathways significantly enriched (Fig. 1D, right panel, adjusted p value < 0.05) at exponential phase in cells grown at 37°C contain genes and/or processes involved in anaerobic respiration, including nitrogen metabolism, porphyrin metabolism, and arginine metabolism (Fig. 2). With those pathway enrichment results in mind, we noticed a striking trend that the genes most highly upregulated at 37°C compared to 25°C (i.e., genes with the highest log_2_FoldChange value in Data Set S1) are those genes involved in anaerobic respiration (22). We cross-referenced genes in Data Set S1 with anaerobic regulation studies (23–37) and found that the majority of known anaerobic genes are upregulated at 37°C compared to 25°C. For convenience, these genes and their corresponding fold changes in thermoregulation are also presented in Table 1 along with if each gene is regulated by Anr, the low oxygen transcriptional regulator. In the absence of sufficient oxygen for fully aerobic respiration, *P. aeruginosa* can utilize various nitrogen oxides as terminal electron receptors for anaerobic respiration in a process known as denitrification (38, 39). The *nar*, *nor*, *nir*, and *nos* operons each encode genes whose products are involved in the reduction of a specific nitrogen oxide and were among the genes most upregulated at 37°C, along with other genes involved in the regulation of denitrification, such as *narXL* and *dnr* (Table 1). Also upregulated at 37°C are *moeA1* and *moaB1*, two genes in the biosynthetic pathway for production of molybdopterin guanine dinucleotide (MGD) cofactor, which is essential for the activity of nitrate reductase and thus anaerobic respiration via denitrification. *P. aeruginosa* contains two operons for cytochrome c *cbb_3_*-type oxidases that accept electrons and reduce oxygen to water. One of the operons, *ccoNOQP*-2, is greatly induced under low oxygen conditions, while the *ccoNOQP*-1 operon is dominant under high oxygen conditions (33). We found all genes of the *ccoNOP*-2 were significantly upregulated at 37°C, while the immediately downstream *ccoNOQP*-1 operon was not thermoregulated (Table 1 and Data Set S1). Under low oxygen and low nitrogen conditions, and thus in the absence of oxygen or nitrogen species as terminal electron receptors, *P. aeruginosa* can survive by converting ADP to ATP via the arginine deaminase pathway encoded by the *arcDABC* operon, which is induced by low oxygen (36, 40–43) and was upregulated at 37°C in our dataset. Both the anaerobic ribonucleoside reductases *nrdD* and *nrdG*, as well as the constitutive ribonucleoside reductases *nrdJ*a and *nrdJ*b were upregulated at 37°C, indicating that warmer temperature may support a faster growth rate by increasing the pool of dNTPs for DNA replication. PAO1 may also experience higher oxidative stress when growing at 37°C compared to 25°C, possibly due to reactive intermediates produced by the denitrification pathway, as catalase *katA* is upregulated (44). Interestingly, as hydrogen cyanide synthesis reactions use electrons to reduce reactive oxygen species to less toxic forms, the *hcnABC* operon may also be indirectly linked to anaerobic respiration; expression of the *hcnABC* is also induced under low oxygen conditions (34). Other metabolic processes were upregulated at 37°C and are shown in Figure 2, with pathways related to survival in low oxygen environments marked as such.

**Figure 2.**
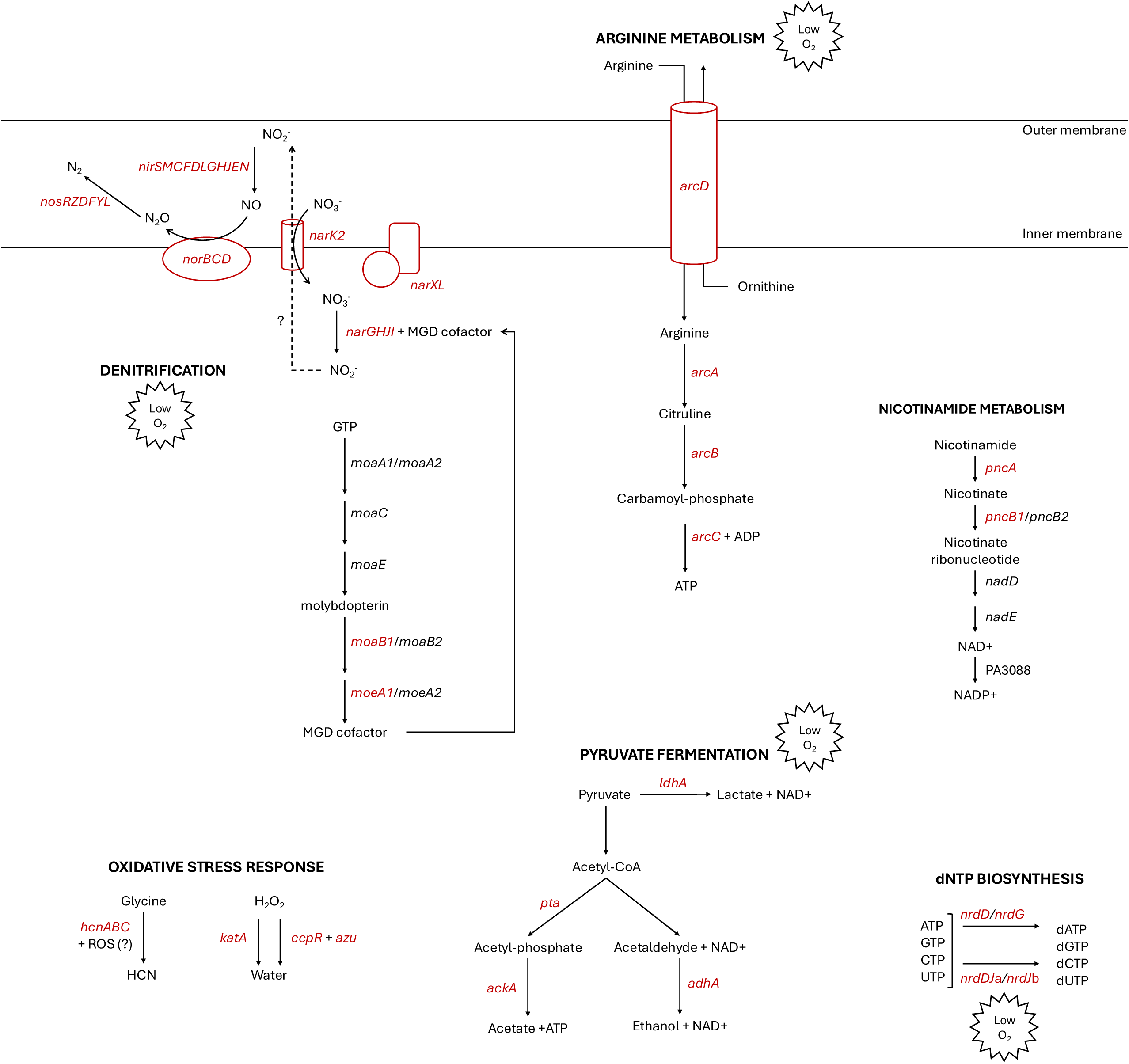
Metabolic pathways upregulated in *P. aeruginosa* growing at 37°C at exponential phase. A selection of notable metabolic and enzymatic pathways that were characteristic of PAO1 growing at 37°C at exponential phase. Genes labeled in red were significantly upregulated at 37°C (fold change > 2, adjusted p-value < 0.05). Pathways are simplified to highlight the function of thermoregulated genes. Pathways or genes related to anaerobic growth and survival and/or induced by low oxygen conditions are marked with Low O_2_. Abbreviations: MGD – molybdopterin guanine dinucleotide; ROS – reactive oxygen species; HCN – hydrogen cyanide,– indicates an incomplete understanding in a specific aspect of a metabolic pathway.

**Table 1.**
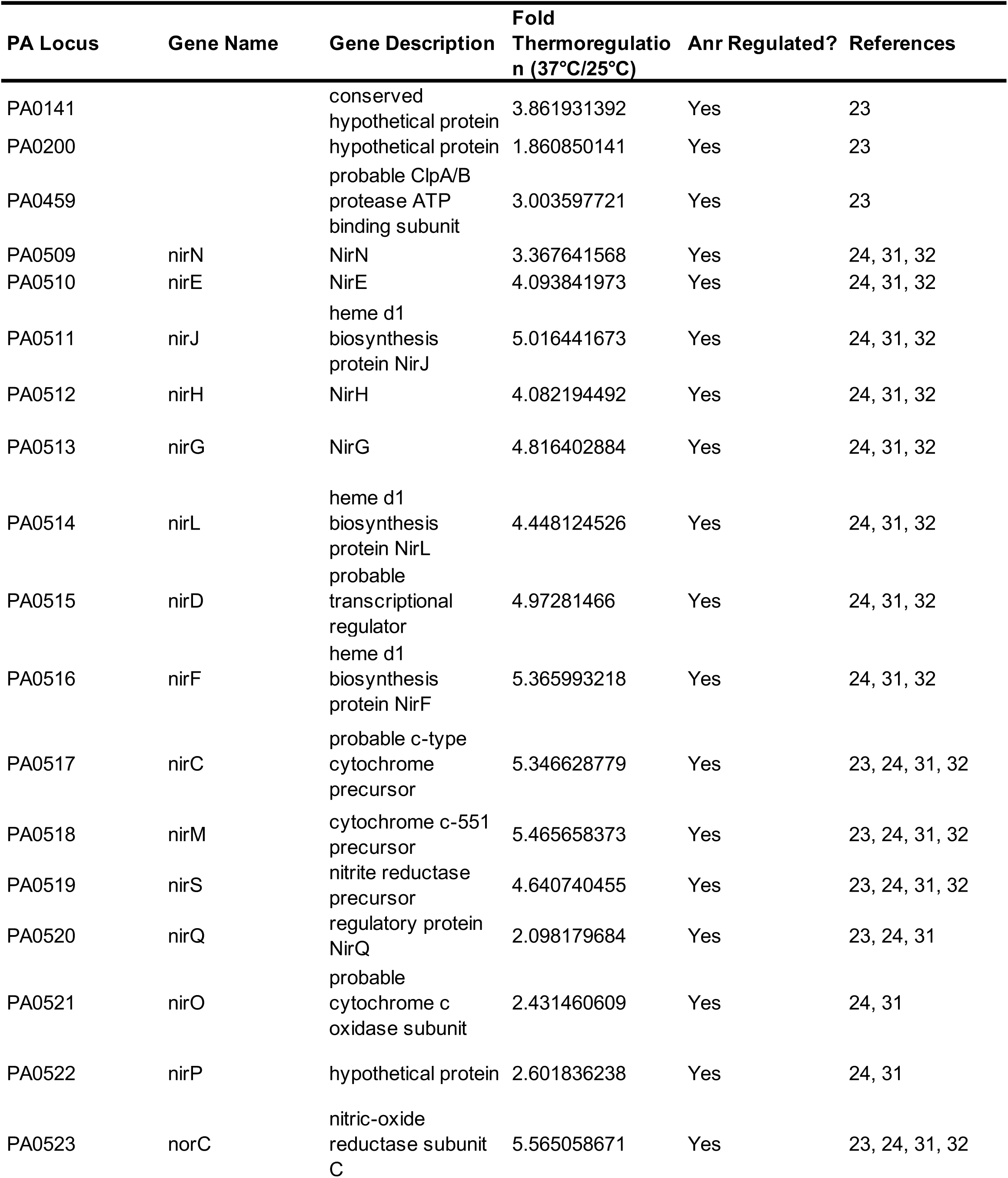
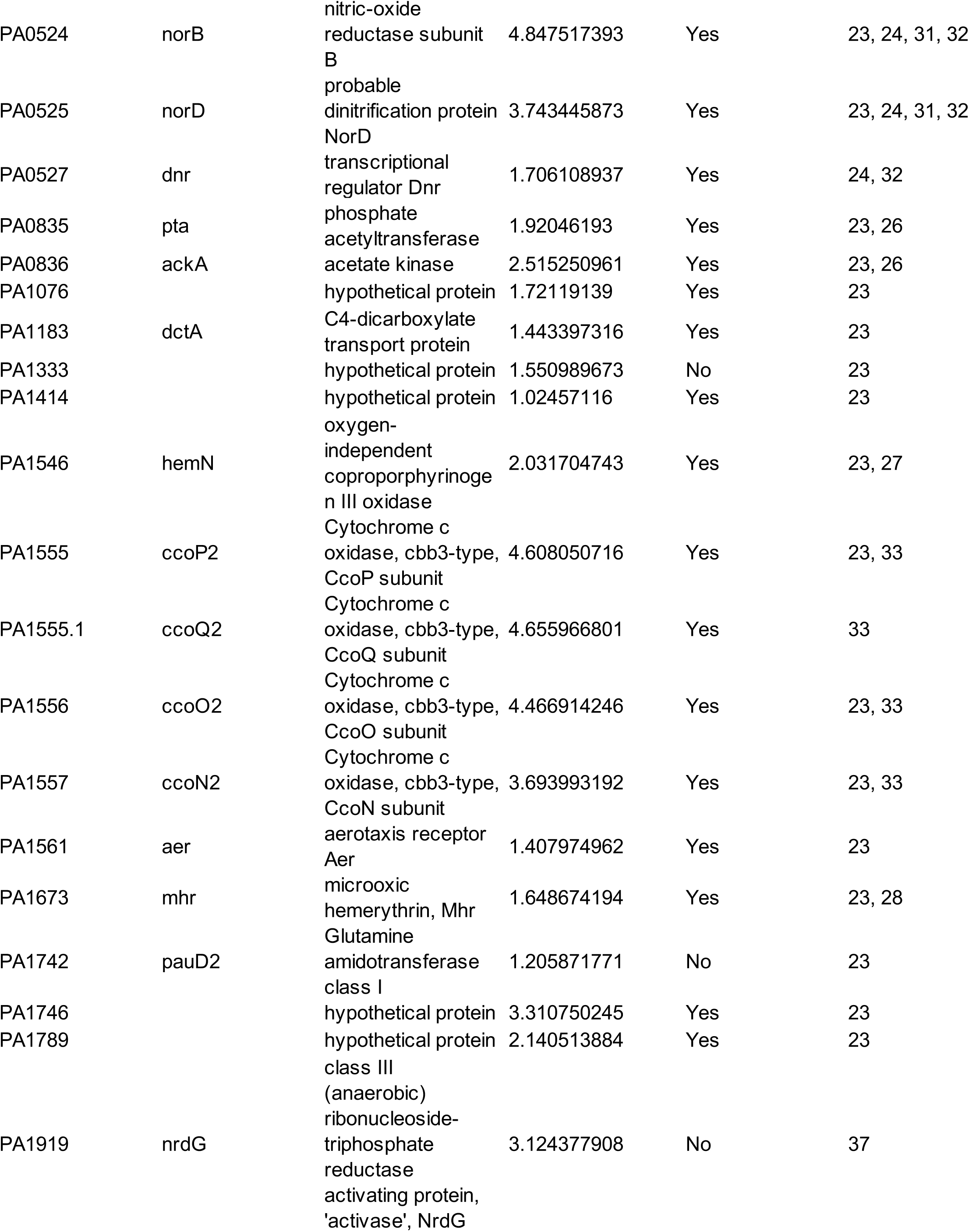
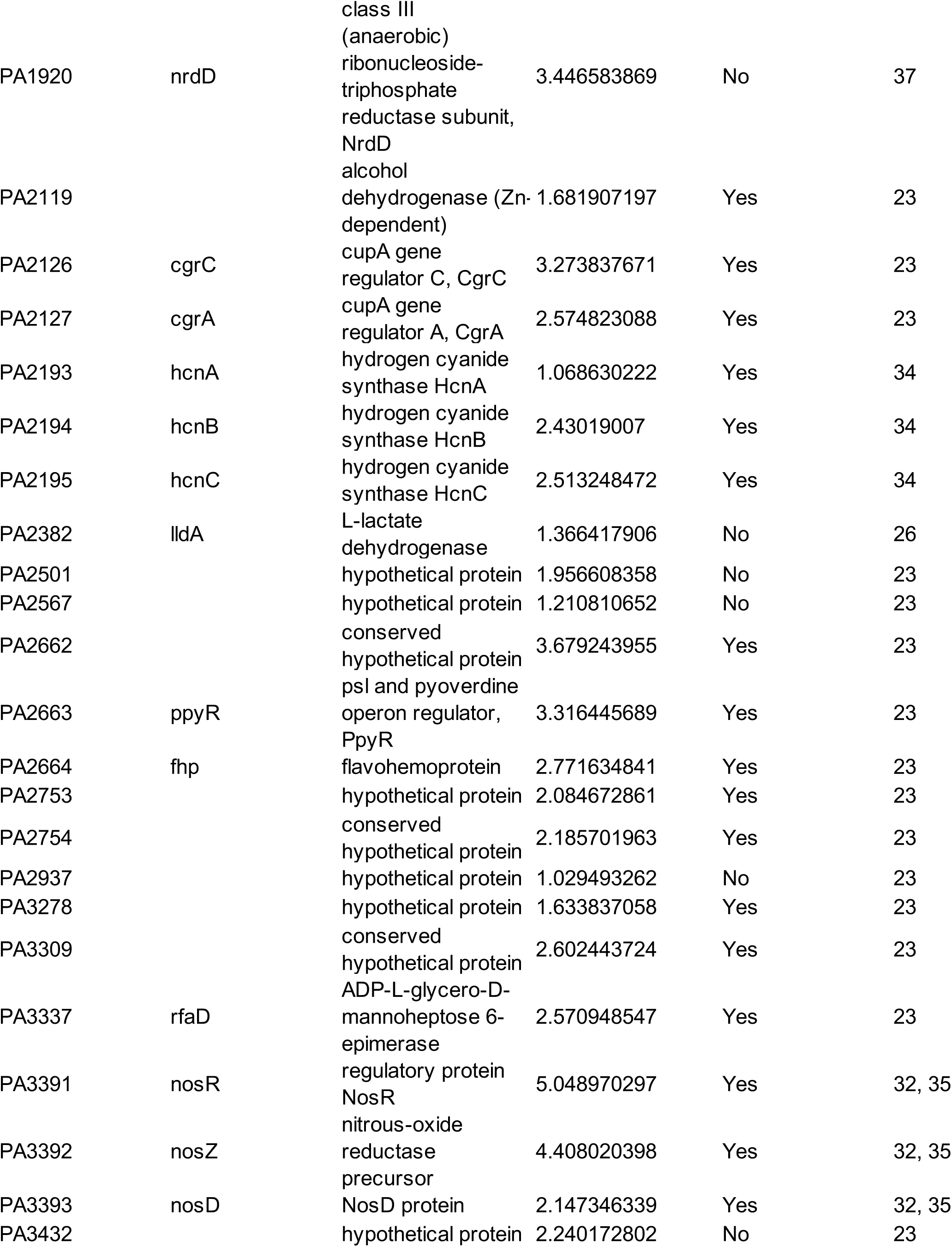
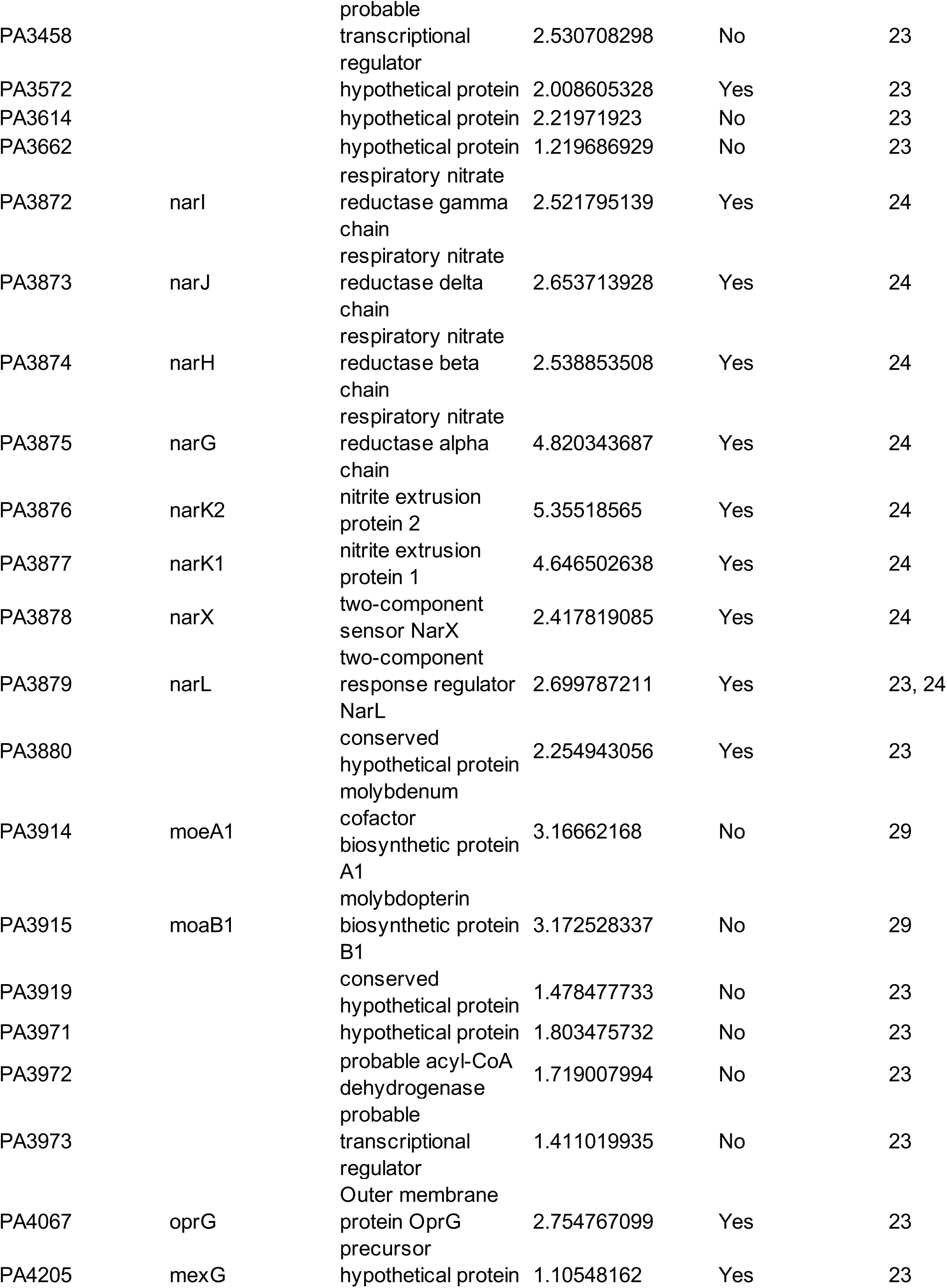
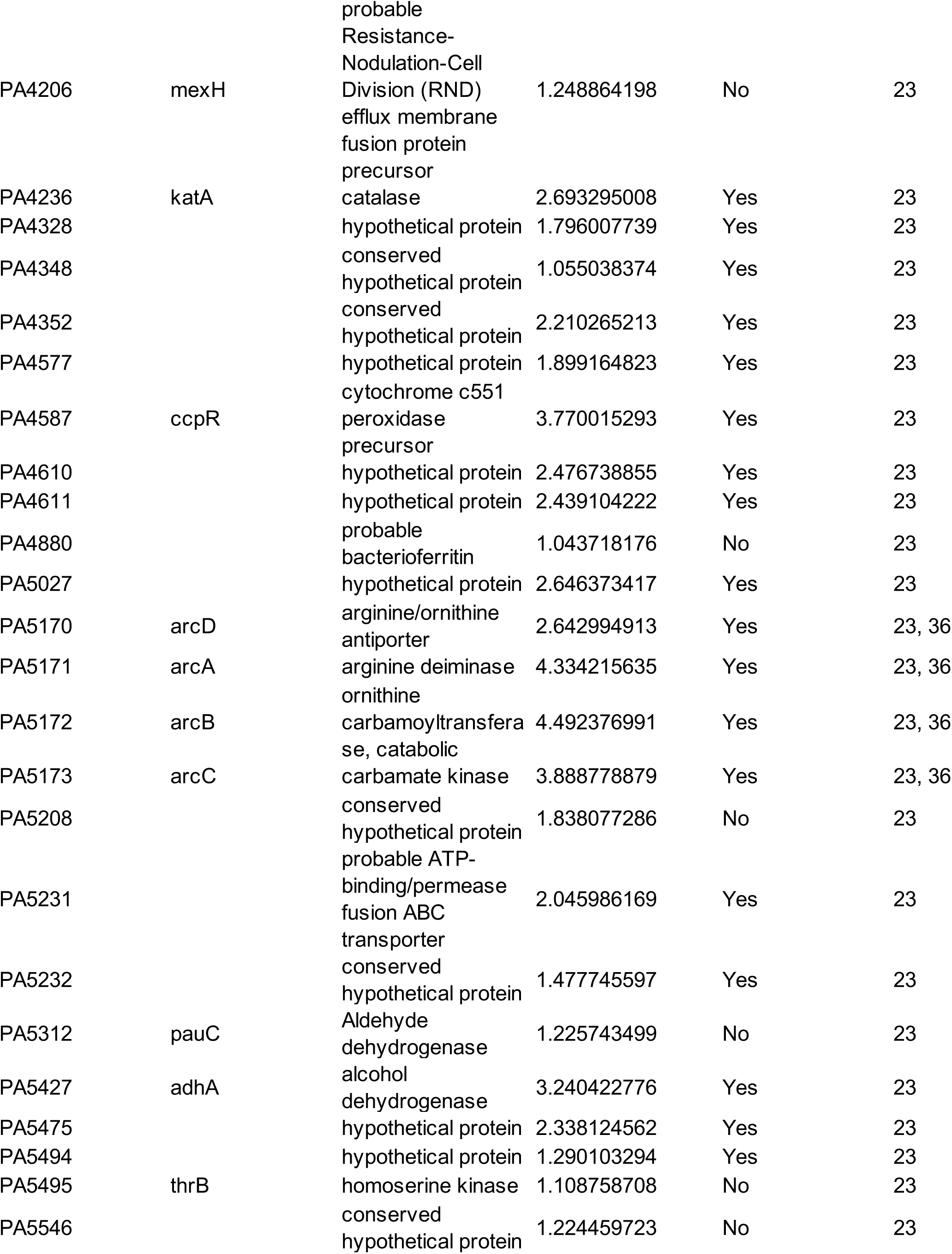
Low oxygen response genes that are thermoregulated.

At 25°C, aminoacyl-tRNA biosynthesis was significantly enriched (Fig. 1D, left panel, adjusted p value < 0.05), which suggests that cells adapt to growing at lower temperatures by increasing tRNA pools for translation. Also upregulated at 25°C were 30S ribosomal subunit genes *rpsU* and *rpsT* (Data Set S1). We initially predicted that cold-shock protein gene(s) would be upregulated at 25°C, as the transition from overnight growth at 37°C to subculturing at 25°C imparts a modest cold shock. We examined the normalized gene expression of all five annotated putative cold-shock proteins in PAO1 (PA0456, PA0961, PA1159, PA2622, PA3266) and found varying thermoregulation and growth phase regulation phenotypes (Fig. 3). PA0456 was expressed equally 25°C and 37°C at both growth phases, and generally higher at exponential phase than stationary phase. PA0961 was lowly or not expressed in any of the conditions tested. PA1159 was also not thermoregulated at either growth phase, although transcript levels appeared slightly higher at stationary phase. PA2622 was lowly expressed at exponential phase and highly expressed at stationary phase, independent of temperature. Only PA3266, also called *capB*, was more highly expressed at 25°C than 37°C at exponential phase (∼6 fold higher, Data Set S1); at stationary phase, expression was low at both 25°C and 37°C and not thermoregulated. CapB was originally identified in the related pseudomonad *Pseudomonas fragi* as one of four low molecular weight proteins induced by cold-shock (45). The authors found that at the amino acid level, CapB was similar to the well-studied *Escherichia coli* cold-shock RNA chaperone CspA (45). Under the conditions we tested, *capB* was the only putative cold-shock protein gene whose expression was induced at 25°C. Its expression greatly decreased by stationary phase, suggesting that *capB* is likely involved in the initial adaptation to colder growth conditions. We also note that *eftM*, a thermolabile methyltransferase that modifies EF-Tu at 25°C but not 37°C in PAO1 which we have previously studied (12), is expressed ∼5.85 fold more at 25°C than 37°C, which is consistent with an enzyme known to be functional only at ambient temperatures (Data Set S1).

**Figure 3.**
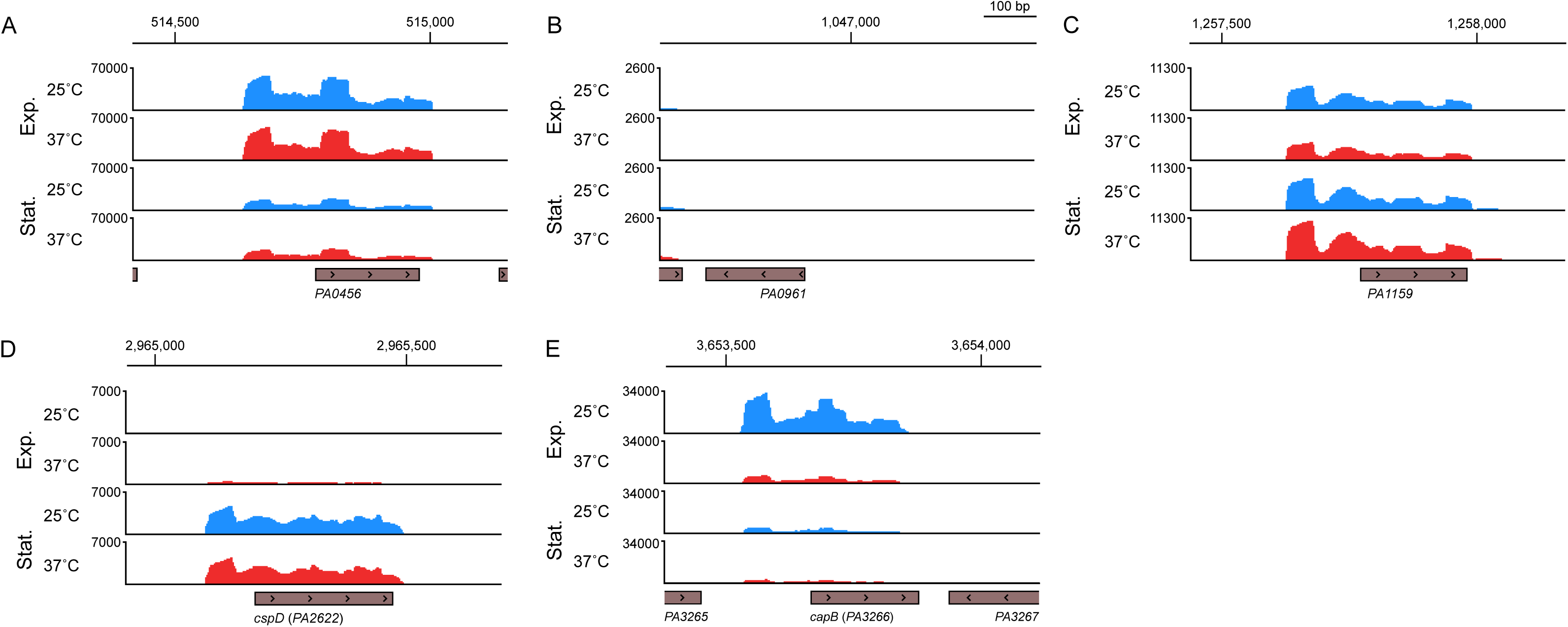
*capB* is the only putative cold shock response gene regulated by temperature. Normalized reads from the RNA-seq experiment diagramed in Fig. 1A are visualized in IGV for the annotated putative cold shock response genes PA0456 (A), PA0961 (B), PA1159 (C), PA2622 (*cspD*) (D), and PA3266 (*capB*) (E). Reads for either the plus or minus strand are shown as relevant to the transcriptional direction of each gene, indicated by arrows within the gray boxes marking the protein coding region of each gene. Reads from the same representative biological replicate sampled first at exponential phase and again at stationary phase are shown for all genes. The genomic position is indicated at the top of each panel.

### Thermoregulation at stationary phase

At stationary phase, expression of 715 genes was affected 2-fold or greater by temperature (adjusted p value < 0.05), with 392 genes upregulated at 37°C compared to 25°C and 323 genes upregulated at 25°C compared to 37°C (Fig. 1C and Data Set S2). At this growth phase, 12.48% of the annotated genome was thermoregulated. This is comparable to our findings for thermoregulation at exponential phase, and a higher percentage than what had been previously found to be thermoregulated at stationary phase in either PAO1 or PA14 laboratory strains (14, 15). We found that phenazine biosynthesis and quorum sensing pathways, including the genes *rhlAB*, *lasA*, and *lecB*, were significantly enriched at 37°C (Fig. 1E, right panel, adjusted p value < 0.05), whereas at 25°C the metabolism of various amino acids as well as glyoxylate metabolism (related to the glyoxylate shunt) were enriched (Fig. 1E, left panel, adjusted p value < 0.05).

To examine how each annotated gene in the genome was thermoregulated at exponential phase versus stationary phase, we compared the fold thermoregulation of each statistically significant gene (adjusted p value < 0.05, a total of 5,018 genes) at exponential phase to its fold thermoregulation at stationary phase (Fig. 1F). Thermoregulation at exponential and stationary phases was weakly correlated (r = 0.1688) and most genes that were thermoregulated were only thermoregulated at one growth phase (Fig. 1G). This underscores that temperature induces transcriptional changes that are distinct to the growth phase of the bacterial population.

To further investigate how temperature and growth phase affect global transcriptional changes, we examined the variation in gene expression from all four conditions studied (exponential at 37°C, exponential at 25°C, stationary at 37°C, stationary at 25°C) using a principal component analysis (Fig. 4A). Despite the different growth temperatures, samples from exponential phase at 25°C and 37°C generally clustered together, as did samples from stationary phase at 25°C and 37°C. Additionally, samples from exponential phase (circles in Fig. 4A) clustered rather distinctly from samples from stationary phase (triangles in Fig. 4A) regardless of temperature, indicating that growth phase contributes more to overall variation than temperature does. Given the distinct clustering based on growth phase, we next used differential expression analysis to compare gene expression at stationary phase versus exponential phase at each 37°C and 25°C. At 37°C, the expression of 2,457 genes (43.00% of the annotated genome) was affected 2-fold or greater by growth phase (adjusted p value < 0.05), with 1,371 genes upregulated at stationary phase and 1,086 genes upregulated at exponential phase (Fig. 4B and Data Set S3). At 25°C, the expression of 2,760 genes (48.31% of the annotated genome) was affected 2-fold or greater (adjusted p value < 0.05) by growth phase, with 1,493 genes upregulated at stationary phase and 1,267 genes upregulated at exponential phase (Fig. 4C and Data Set S4). We then compared the growth phase regulation of each statistically significant gene (adjusted p value < 0.05, a total of 5,018 genes) at 37°C to its grown phase regulation at 25°C (Fig. 4D) and found a positive correlation between growth phase regulation at 37°C and 25°C (r = 0.8637). While most genes were regulated by growth phase similarly at both 37°C and 25°C, there were some exceptions. The *glc* operon genes were only upregulated at stationary phase when cells were grown at 25°C but not when grown at 37°C (Fig. 4D and Data Sets S3 and S4). In conclusion, growth phase regulates gene expression similarly whether PAO1 is grown at 37°C or 25°C and is a stronger environmental cue driving global transcriptional changes.

**Figure 4.**
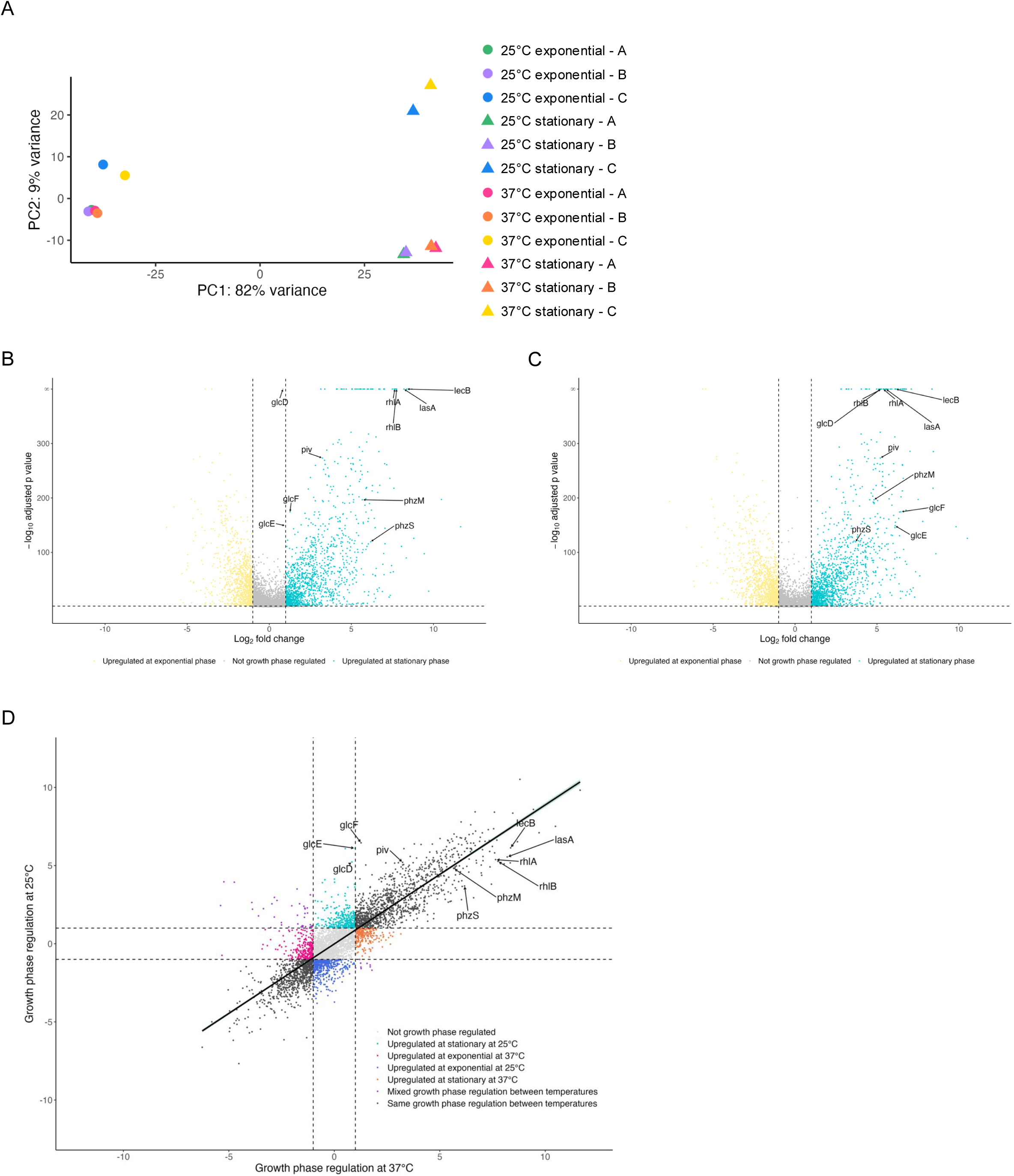
Growth phase regulates genes similarly at 25°C and 37°C. A) Principal component analysis of reads from RNA-seq of three biological replicates of PAO1 grown at two temperatures and sampled at two growth phases. Each data point represents a transcriptome of PAO1 grown at 25°C (cool colors) or 37°C (warm colors) from exponential phase (circles) or stationary phase (triangles). Each color indicates a specific bacterial population that was sampled first at exponential phase and again at stationary phase to allow longitudinal comparison. B, C) Volcano plots showing the growth phase regulation of PAO1 transcripts at 37°C (B) and 25°C (C). Differential gene expression analysis (DESeq2) was used to compare gene expression at stationary phase to exponential phase. Transcripts with an absolute value of fold change greater than 2 (vertical dashed lines) and an adjusted p-value < 0.05 (horizontal dashed line) were considered growth phase regulated, with transcripts upregulated at stationary phase represented by teal points and transcripts upregulated at exponential phase represented by yellow points. Transcripts that did not change by an absolute value of fold change greater than 2 or were not statistically significant (adjust p-value > 0.05) are depicted in gray. Genes of interest are annotated. D) The fold growth phase regulation (expression at stationary/exponential phase) of statistically significant transcripts was plotted at 37°C (x-axis) versus 25°C (y-axis). Transcripts are colored according to how temperature affected growth phase regulation and genes of interest annotated. Linear regression with 95% confidence intervals is shown.

### LasR regulation is generally robust against temperature changes

We noticed that many genes known to be regulated by LasR were upregulated at 37°C compared to 25°C at stationary phase (Data Set S2). As with many transcriptional regulators, LasRI quorum sensing regulation has been historically studied at 37°C (16–19, 46). With our recent finding that the secreted protease *piv*, which is upregulated at 25°C compared to 37°C at stationary phase, is regulated by LasR in a temperature-dependent manner (9), we wondered if LasR could regulate other genes in a temperature-dependent manner. To address this, we grew PAO1 and an isogenic Δ*lasR* mutant overnight at 37°C and then subcultured in biological triplicate at 37°C or 25°C (Fig. 5A). At stationary phase (OD_600_ of 2.0), when LasR regulation is active, RNA was extracted and RNA-sequencing was performed. We first determined genes regulated by LasR at 37°C and found that the expression of 778 genes was changed 2-fold or greater (adjusted p value < 0.05) in the Δ*lasR* mutant (Fig. 5B and Data Set S5). Of those genes, the expression of 503 genes decreased in Δ*lasR* while the expression of 275 genes increased. At 25°C, the expression of 667 genes was changed 2-fold or greater (adjusted p value < 0.05) in the Δ*lasR* mutant (Fig. 5C and Data Set S6). Of those, the expression of 424 genes decreased in Δ*lasR* while the expression of 243 genes increased. These data reveal that LasR regulates similar numbers of genes at 37°C as at 25°C, both positively and negatively.

**Figure 5.**
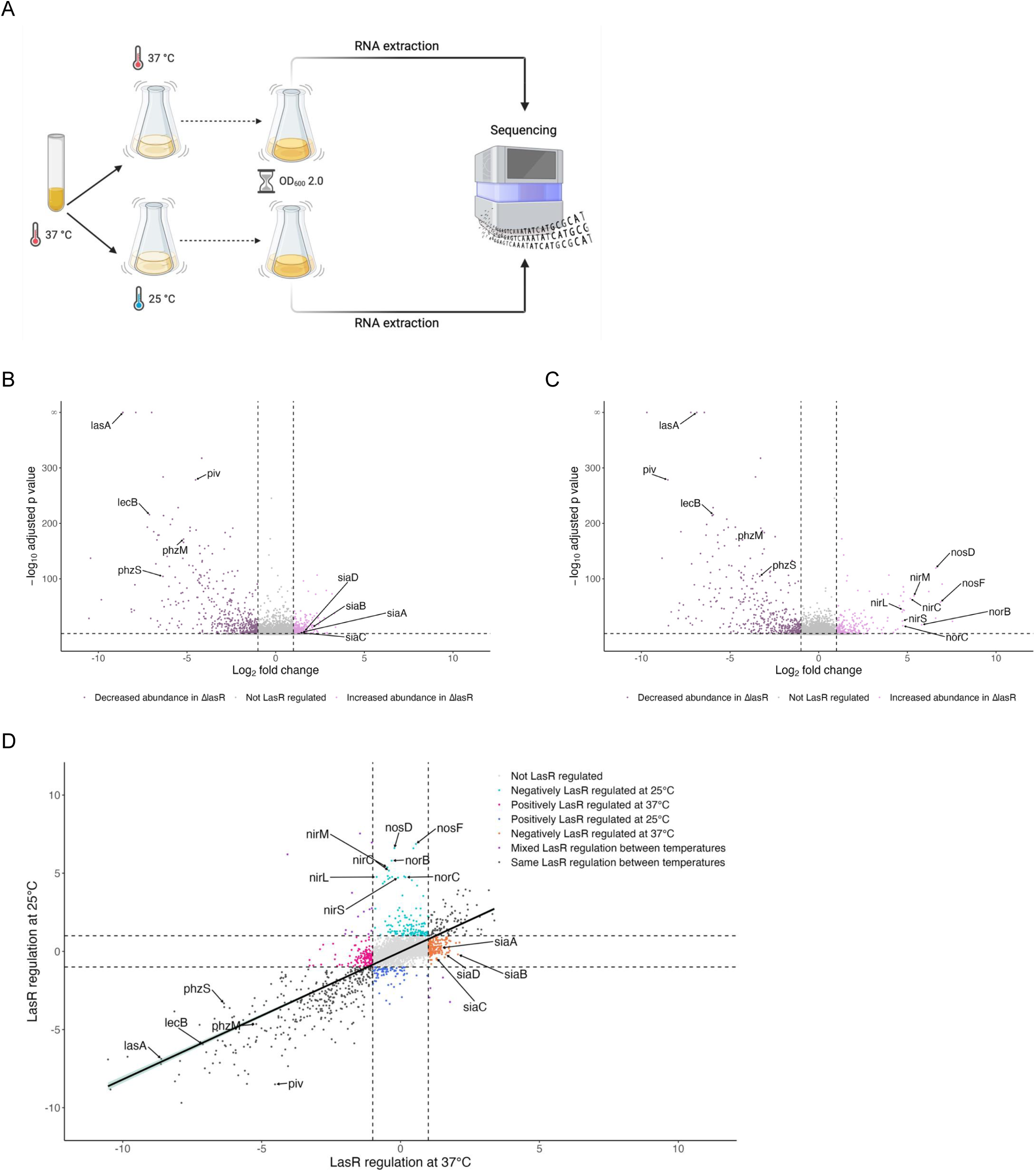
LasR regulates most target genes similarly at 25°C as 37°C, with notable exceptions. A) Diagram of experimental setup for RNA-seq of PAO1 and Δ*lasR* grown at two temperatures and sampled at stationary phase. PAO1 and Δ*lasR* were grown overnight at 37°C and then subcultured in parallel at 37°C and 25°C. At stationary phase, RNA was extracted from ∼10^9^ cells and sequenced on an Illumina NovaSeq X Plus. Created using BioRender.com. B, C) Volcano plots showing the LasR regulation of PAO1 transcripts at 37°C (B) and 25°C (C). Differential gene expression analysis (DESeq2) was used to compare gene expression in Δ*lasR* to PAO1. Transcripts with an absolute value of fold change greater than 2 (vertical dashed lines) and an adjusted p-value < 0.05 (horizontal dashed line) were considered LasR regulated, with transcripts negatively regulated by LasR represented by light pink points and transcripts positively regulated by LasR represented by purple points. Transcripts that did not change by an absolute value of fold change greater than 2 or were not statistically significant (adjust p-value > 0.05) are depicted in gray. Genes of interest are annotated. D) The fold LasR regulation (expression in Δ*lasR*/PAO1) of statistically significant transcripts was plotted at 37°C (x-axis) versus 25°C (y-axis). Transcripts are colored according to how temperature affected LasR regulation and genes of interest annotated. Linear regression with 95% confidence intervals is shown.

To investigate how temperature affects LasR regulation, we compared the LasR-regulation of each statistically significant gene (adjusted p value < 0.05, a total of 3,402 genes), including those not differentially expressed, at 37°C with its LasR-regulation at 25°C (Fig. 5D). We found a positive correlation between LasR regulation at 37°C and 25°C (r = 0.776), indicating that LasR generally regulates target genes similarly at both 37°C and 25°C, including the well-studied LasR targets *lasA*, *lecB*, and *rsaL*. We also found that LasR regulated *piv* more at 25°C than 37°C, an observation consistent with our previous findings (9). To identify those genes that were not regulated by LasR similarly at both temperatures (Table S1), we then further filtered these genes based on their LasR-regulation at 37°C and 25°C according to the following criteria:

a) a gene was regulated by LasR at one temperature but not the other (i.e., a gene was differentially expressed in Δ*lasR* at only one temperature) or
b) a gene was regulated by LasR positively (or negatively) at one temperature and vice versa) at the other temperature.

Interestingly, the LasR-regulation of over 500 genes was sensitive to temperature and note that the majority met the first criteria of being regulated by LasR at only one temperature and hence are present in Data Set S5 *or* S6, but not both. Notably, the *nirNEJHGHLDFCMS*, *nirQOP*, *norBC*, and *nosRZDF* operons were strongly upregulated in Δ*lasR* compared to PAO1 at 25°C only (Table S1). This indicates that normally in PAO1 cultures at 25°C, LasR negatively regulates the *nir*, *nor*, and *nos* operons, and, at 37°C, LasR does not regulate these operons. Of the nitrate reductase *nar* operon, only two genes (*narK1* and *narH*) had their LasR regulation affected by temperature, which suggests that the temperature-dependent LasR regulation of denitrification at 25°C may be limited to the *nir*, *nor*, and *nos* operons. We also found that the *siaABCD* operon, which promotes biofilm formation in response to SDS exposure and/or carbon availability (47, 48), was negatively regulated by LasR at 37°C (i.e., *siaABCD* expression was increased in Δ*lasR* compared to PAO1) but not at 25°C.

## DISCUSSION

Using RNA-seq, we identified the global thermoregulon for PAO1 at exponential phase (Data Set S1) and stationary phase (Data Set S2), as well the growth phase regulon at 37°C (Data Set S3) and at 25°C (Data Set S4). We also determined the LasR regulon at stationary phase using a clean Δ*lasR* mutant, at both the commonly studied temperature of 37°C (Data Set S5) as well as the more environmentally relevant temperature of 25°C (Data Set S6). Our experimental design for paired growth at two temperatures with longitudinal sampling at two growth phases allowed us to compare temperature and growth phase as two environmental factors driving the transcriptomic landscape. We found that ∼13% of the PAO1 transcriptome was thermoregulated at each exponential and stationary phase, which is higher than prior studies that found ∼6% thermoregulated at only stationary phase in *P. aeruginosa* strains PAO1 (14) and PA14 (15). This could be due to differences in strains used (PAO1 vs PA14), differences in sequencing technologies used (microarray vs RNA-sequencing), better annotation of the PAO1 genome, or other technical aspects such as the improvement of RNA-sequencing technology and methods of downstream analysis. Distinct genes were thermoregulated at each growth phase, largely due to genes being expressed at one growth phase but not the other. With exceptions (such as the *glc* operon, which is only growth phase regulated at 25°C), growth phase affected gene expression similarly at 25°C and 37°C (Fig. 4D). Growth phase also affected more of the transcriptome than temperature, with upwards of 50% of the transcriptome regulated by growth phase at each 25°C and 37°C, and transcriptomes from the same growth phases were more similar to each other regardless of the temperature conditions (Fig. 4A). This underscores the role of growth phase and kin cell density in *P. aeruginosa* physiology and the inherent robustness of *P. aeruginosa* to grow at different temperatures without extensive changes to the transcriptome. Accordingly, we found that expression of many of its oft studied virulence factors and behaviors, such as the genes for rhamnolipid, lectin, and pyocyanin production, is more sensitive to growth phase than temperature changes. This indicates that although *P. aeruginosa* clearly adapts to growing at different temperatures, upregulation of its key virulence factors is largely unaffected by the perturbations in temperature that we tested and are expressed even at non-human body temperature. This supports the hypothesis that these virulence factors are general mechanisms for this bacterium to acquire nutrients and compete with other microbes, and also happen to be beneficial for survival in humans by facilitating pathogenesis.

A major question of this work is how *P. aeruginosa* responds to growing at different temperatures at exponential phase and stationary phase. At exponential phase, we found that increased expression of many genes related to anaerobic growth and survival (22) is a strong signature of growth at 37°C versus 25°C (Fig. 1D and Table 1.) This included both genes for anaerobic respiration via denitrification (*nar*, *nir*, *nor*, and *nos* operons) as well as non-nitrogen-based mechanisms of survival in low oxygen environment (the high oxygen affinity type II cytochrome c oxidase *ccoNOQP*-2, the arginine deaminase pathway *arcDABC*, pyruvate fermentation, and others, see Fig. 2 and Table 1). Of note is that many of these thermoregulated genes related to anaerobic survival are regulated by Anr (Table 1), a major low oxygen transcriptional regulator in *P. aeruginosa* (25), and thus we suspect that regulation by Anr could be affected by temperature. However, expression of the *anr* gene is not thermoregulated in our datasets and thus does not explain how virtually all of the known Anr regulon is thermoregulated. As our study was limited to RNA-sequencing, there could be post-transcriptional or translational mechanisms of thermoregulation acting on Anr to alter its regulatory activities in a temperature-dependent manner. Regulation by Anr is complex and has been highly studied (22, 39); in addition to its direct targets, Anr also upregulates the regulator Dnr, which specifically activates the *nar*, *nir*, *nor*, and *nos* denitrification operons and is important for fitness in microoxic as well as anaerobic environments (49). Expression of *dnr* was slightly thermoregulated in our dataset (Table 1 and Data Set S1), which could contribute in part to the thermoregulation of its specific targets. However, since much of the Anr regulon is not co-regulated by Dnr, thermoregulation of *dnr* cannot fully account for how so many anaerobic genes are also regulated by temperature and future studies are needed.

Although we studied gene expression in *in vitro* lab conditions, we also note that many of the oxygen starvation genes upregulated at 37°C during exponential growth are upregulated in respiratory infections in CF patients (50, 51). Mucus within the lungs of CF patients has been characterized as both oxygen-poor and sufficiently nitrogen-rich for *P. aeruginosa* to respire via denitrification (52–56). That temperature alone can lead to upregulation of oxygen starvation genes associated with chronic infection and long-term adaptation to the CF lung environment (22, 50, 51) suggests that thermoregulation may function to prime *P. aeruginosa* for adaptation to the human body as a new environment. This work thus provides new insights into how thermoregulation participates in *P. aeruginosa*’s successful transition from an ambient environment to the human body.

For genes upregulated at exponential phase at 25°C, we expected to identify cold-shock response genes, as transitioning an overnight culture grown at 37°C to subculturing at 25°C simulates a mild cold shock that slows the growth of PAO1, particularly during the initial lag and early exponential phases of growth (9). However, we found that expression of only one of the five annotated putative cold shock proteins, PA3266 (*capB*), was upregulated at 25°C, and the

remaining four were either not expressed under the tested growth conditions or were expressed during at least one growth phase but were not thermoregulated (Fig. 3). Cold-shock proteins are commonly identified by a characteristic and highly conserved nucleotide binding domain of an anti-parallel, five strand β-barrel first identified in eukaryotic Y-box proteins (57). CapB appears similar to the well-studied cold-shock RNA binding protein CspA in *Escherichia coli*; in *E. coli*, CspA serves as a ‘master’ cold shock regulator and negatively regulates other cold-shock genes such that they are only expressed if CspA becomes non-functional (58–60). Thus, the other annotated putative cold shock genes may be similarly repressed by CapB in *P. aeruginosa*, or they may respond to other environmental stressors than temperature, as is the case for CspD in *E. coli* being induced by nutritional deprivation at stationary phase (61); we also found that expression of PA2622, annotated as CspD in the PAO1 genome, was induced at stationary phase at both 25°C and 37°C.

At stationary phase, we noted many thermoregulated genes are also LasR-regulated, which has been previously observed (14, 15) and was not unexpected as LasR is an active regulator at stationary phase and drives many (although not all) transcriptional changes during this growth phase. The LasRI quorum sensing regulon has previously been characterized by adding the autoinducer 3O-C12-HSL to a *P. aeruginosa* strain deficient in producing any quorum sensing signal (16, 18, 19) or by ectopically overexpressing the LasR receptor during early growth in addition to supplementation with 3O-C12-HSL. The effects of temperature on LasR regulation had not been previously explored and we thus determined the LasRI quorum sensing regulon using a clean Δ*lasR* mutant at both 37°C and 25°C at stationary phase, when LasR would be active in wild-type cells. Overall, LasR regulates many of its target genes to a similar degree when grown at either 37°C or 25°C (Fig. 5D). However, we found over 500 genes whose LasR regulation is sensitive to temperature (Table S1), including genes that have not previously been recognized as LasR-regulated, possibly due to all prior transcriptomic studies of the LasR regulon being conducted at 37°C (17–19, 46). One example of this is the *nir*, *nor*, and *nos* operons, which are negatively regulated by LasR at 25°C but not regulated by LasR at 37°C (see Data Sets S6 and S5, respectively) and have not been previously found to be regulated by LasR specifically. One study of a strain deficient in both LasRI and RhlRI systems has found that quorum sensing in general regulated the *nar*, *nir*, *nor*, and *nos* operons (16), although this regulation could not be attributed specifically to LasR. Furthermore, subsequent studies of the LasR specific regulon failed to identify genes in the *nir*, *nor*, and *nos* operons (17, 18, 46). As LasR is known to function only as a direct positive regulator, we speculate that negative regulation of the *nir*, *nor*, and *nos* operons occurs through LasR activating a repressor only at 25°C or activating a repressor that itself is functional at 25°C but not 37°C. Previous studies conducted at 37°C have also found genes repressed by the LasRI quorum sensing system despite LasR being a direct transcriptional activator (16, 18). Growing *P. aeruginosa* at the non-standard temperature of 25°C was critical for finding these previously unrecognized LasR-regulated genes and these data underscore the importance of diverse environmental and nutritional conditions in better understanding bacterial physiology.

In conclusion, as an opportunistic human pathogen, *P. aeruginosa* is capable of surviving in both an ambient environment and the human body and must sense the transition between these environments in order to adapt accordingly. Studying the transcriptome at an ambient temperature, 25°C, as well as human body temperature, 37°C, at both exponential and stationary phase has led to new insights in the role of thermoregulation in *P. aeruginosa* adapting to the human body. Our RNA-seq analyses have also identified many thermoregulated genes for future research on the mechanism(s) by which temperature regulates their expression.

## MATERIALS AND METHODS

### Culture conditions and RNA-sequencing

Biological triplicates of the indicated strain (Table 2) were grown in 3 mL lysogeny broth (LB) overnight in a rolling drum at 37°C,subcultured to an initial OD_600_ of 0.05 in 25 mL LB, and incubated at either 25°C or 37°C with shaking at 200 rpm. RNA was extracted from ∼10^9^ cells of the triplicate cultures first at exponential phase (OD_600_ = 0.5) and then from the same cultures again at early stationary phase (OD_600_ = 2.0) using TRI-Reagent (Millipore Sigma) according to the manufacturer’s recommendations. For the RNA-seq comparing PAO1 and Δ*lasR* at 25°C and 37°C, RNA was extracted from biological triplicate cultures at early stationary phase. All samples were treated with TURBO DNase (ThermoFisher) and RNA-seq subsequently performed by SeqCenter (Pittsburgh, PA) on an Illumina NextSeq2000 or NovaSeq X Plus as indicated. Libraries were prepared by SeqCenter using Illumina Stranded Total RNA Prep Ligation with Ribo-Zero Plus kit for rRNA depletion.

**Table 2.** Bacterial strains used in this study.

| Strain | Source |
| --- | --- |
| *Pseudomonas aeruginosa* PAO1 | Simon Dove (Boston Children's Hospital) |
| PAO1 Δ*lasR* | Robinson et al., 2025 (9) |

### Analysis of RNA-sequencing

Demultiplexing, quality control, and adapter trimming was performed by SeqCenter using bcl-convert. Reads were mapped to the *Pseudomonas aeruginosa* PAO1 reference genome (NCBI Reference Sequence NC_002516.2) using bowtie2 version 2.4.5 on the following settings: very-sensitive, non-deterministic, dovetail, no-mixed, no-discordant, no-unaligned (62). Mapped reads were counted with htseq-count version 2.0.2, differential gene expression analysis conducted using DESeq2 version 1.38.3 (63), and visualized with volcano plots made using ggplot2. For principal component analyses (PCA), mapped reads were variance stabilizing transformation (VST) normalized using the *vst* function of DESeq2 and subsequently used by the *plotPCA* function of DESeq2 to produce the PCA, which was further visualized using ggplot2. Gene set enrichment analyses of KEGG pathways (in Fig.1D, E) was conducted using the *gseKEGG* function of the clusterProfiler version 4.10.1 and visualized using the *dotplot* function of enrichplot version 1.22.0. Mapped reads were normalized and visualized (in Fig. 3) in Integrative Genomics Viewer (IGV) (64). Additional information on gene names, descriptions, and locus numbers was sourced from the *Pseudomonas* Genome Database (65).

Sequencing data have been deposited to the Gene Expression Omnibus (GEO) at NCBI under accession number GSE304330 and are freely available.

## ACKNOWLEDGEMENTS

We thank the Goldberg Lab for discussion and feedback on the manuscript. R.E.R. was supported by an NIH NRSA F31 Pre-Doctoral Fellowship AI172335. M.J.G was supported by NIH grant GM156848. This study was supported in part by the Emory Integrated Genomics Core (EIGC), which is subsidized by the Emory University School of Medicine and is one of the Emory Integrated Core Facilities. RNA-seq libraries were constructed and sequenced at SeqCenter.

